# Plasmidome, resistome, and virulence-associated genes characterization of *Acinetobacter johnsonii* in NASA cleanrooms and a clinical setting

**DOI:** 10.1101/2025.10.09.681420

**Authors:** Anna Tumeo, Georgios Miliotis, Andy O’Connor, Varsha Vijayakumar, Pratyay Sengupta, Francesca McDonagh, Aneta Kovarova, Christina Clarke, Brigid Hooban, Nitin Kumar Singh, Alexandre Soares Rosado, Karthik Raman, Kasthuri Venkateswaran

## Abstract

Evidence shows persistence of non-spore-forming *Acinetobacter johnsonii* in high-stakes controlled and nutrient-limited environments. This study aims to explore the mechanisms underpinning such adaptability through a comprehensive genomic analysis of 22 isolates of *A. johnsonii* from NASA’s Payload Hazardous Servicing Facility (PHSF) and one carbapenem-resistant strain (E154408A) from patient colonization in Ireland. Core-genome phylogeny revealed clustering of PHSF-originating isolates in a monophyletic clade divergent from the main species lineage. Species-wide virulence-associated genes and metabolic profiling indicated the unique presence in PHSF-originating isolates of two complete efflux pumps and of a conserved allantoin racemase, suggesting adaptability for multiple environmental stresses. Observed ubiquity of *bla*_OXA_ in investigated genomes (n=112) and phenotypically-validated multidrug-resistant profile of E154408A strain highlight *A. johnsonii*’s potential as antimicrobial resistance (AMR) reservoir. Plasmidome analysis suggested gain/loss events across the monophyletic population and potential AMR acquisition pathways. Genome-to-metagenome mapping identified genomic signatures of *A. johnsonii* in PHSF >10 years post initial isolation.

**Importance:** *Acinetobacter johnsonii* is increasingly recognized as an emerging human pathogen, with growing evidence of its ability to persist in controlled, high-stakes environments, posing risks as both persisting environmental contaminant and antimicrobial resistance (AMR) reservoir. Yet, gaps remain in our understanding of its AMR profile and the mechanisms that enable its enhanced environmental adaptability. This knowledge is necessary in contexts where biological cleanliness is a priority such as clinical settings and spacecraft assembly facilities cleanrooms, where contamination of hardware with terrestrial microorganisms is concerning. In this study, we aim to address some of key knowledge gaps by providing genomic insights into a rare multi-drug resistant clinical isolate and 22 NASA cleanroom isolates that persisted for over a decade in extremely clean conditions. Our findings will help evaluate the contamination risk of *A. johnsonii* in high-stakes environments and ultimately strengthen our ability to manage this microbial contaminant across terrestrial and extraterrestrial settings.

**Highlights:**

- Cleanrooms-derived *A. johnsonii* genomes show favorable traits for increased adaptability
- Genomic signatures of *A. johnsonii* persisted in the cleanrooms for >10 years
- *bla*_OXA_ is ubiquitously found in the genome of all *A. johnsonii*
- E154408A is the first patient colonization by carbapenem-resistant *A. johnsonii* in Europe

## Introduction

The *Acinetobacter* genus is known to thrive in controlled environments, largely due to robust biofilm formation enabling survival of desiccation and nutrient-limited conditions (1) and resistance to cleaning and disinfection efforts (2). In addition, the widespread presence of efflux pumps (e.g., AdeABC) in the genomes of *Acinetobacter* species as well as their known tendency to acquire antimicrobial resistance genes (ARGs) to clinically relevant antibiotics (e.g., *bla*_OXA_ beta-lactamases) (3) facilitate their adaptability for multiple environmental stresses and their persistence despite exposure to heavy metals and antibiotics residues (4). *Acinetobacter* species presenting increasingly complex Antimicrobial Resistance (AMR) profiles therefore pose significant clinical challenges when associated with hospital-acquired infections.

*Acinetobacter johnsonii* is a Gram-negative, non-spore-forming, non-fermentative coccobacillus within the *Acinetobacter* genus (5). As a species it exhibits ecological versatility, commonly isolated from environmental niches such as agricultural soil, freshwater systems including rivers and lakes, marine ecosystems, and anthropogenically impacted sites. Although less understood compared to *A. baumannii*, the clinical relevance of *A. johnsonii* is also recently gaining attention. *A. johnsonii*‘s adaptability to desiccation and low-nutrient conditions has indeed been highlighted by its isolation from clinical settings (6) and NASA Spacecraft Assembly Facilities (SAF) cleanrooms (7). A 2020 phylogeographical study identified *A. johnsonii* isolates from both environmental and clinical sources circulate globally and harbor antibiotic-resistance genes including carbapenemases, underscoring its clinical relevance (8). *A. johnsonii* has also been implicated in opportunistic infections such as bacteremia, meningitis, and post-traumatic wound infections, often in immunocompromised hosts (9). Furthermore, scarce reports of carbapenemase producing *A. johnsonii* isolates carrying *bla*_OXA_ and *bla*_NDM-1_ highlight its potential as understudied emerging AMR threat (10).

Critical research and knowledge gaps remain regarding the genomic adaptations underpinning not only *A. johnsonii*’s role as potential reservoir or vector of AMR within healthcare settings but also its persistence as non-spore-forming microorganism in harsh and nutrient-limited environments and throughout enduring disinfection protocols, such as those carried out within SAF cleanrooms. Ensuring biological cleanliness while assembling and launching spacecraft is, however, critical in the search for life elsewhere in the Solar System, and contamination of hardware with terrestrial microorganisms is concerning (7).

In this study, we aim to address some of these knowledge gaps by conducting a comprehensive genomic analysis of 22 *A. johnsonii* isolates, isolated during and after the assembly and testing of NASA’s Mars Phoenix lander at the Payload Hazardous Servicing Facility (PHSF) in the Kennedy Space Center (KSC, Florida, USA), where they survived rigorous disinfection protocols, and of strain E154408A, documented as the first reported carbapenem-resistant patient colonization case in Ireland and Europe. Alongside species-wide genomic characterization, we investigate the AMR profiles of PHSF-derived isolates and E154408A strain and validate their resistance to clinically relevant antibiotics with phenotypic assays. Our findings will help evaluate the contamination risk of *A. johnsonii* in high-stakes sterile environments, inform the development of effective mitigation and antimicrobial stewardship strategies, and ultimately strengthen our ability to manage *A. johnsonii* across both terrestrial and extraterrestrial settings.

## Methods

### Samples collection

22 isolates of *A. johnsonii* were recovered from cleanroom environmental samples collected at KSC-PHSF both during (2P; June 27, 2007) the assembly and testing operations of the NASA Mars Phoenix lander (n=5) and immediately after (3P; August 1, 2007) Phoenix was transferred to the launch pad (n=17). One additional clinical multidrug-resistant (MDR) isolate of *A. johnsonii* (E154408A strain) was isolated in March 2019 through a rectal swab from a 73 years-old patient at Galway University Hospital during routine screening for carbapenem-resistant organisms by Galway Reference Laboratory Service. The patient did not show any signs of infection.

### DNA sequencing and genome assembly

PHSF-originating *A. johnsonii* isolates (n=22) were sequenced using third generation DNA sequencing (Oxford Nanopore Technologies, Oxford, UK) with FLO-PRO114M, R10.4.1 technology. Raw reads were quality controlled using <u>FastQC</u> v0.12. Genomic DNA assembly was conducted using <u>Canu</u> (11) v2.2 and <u>Flye</u> (12) v2.9.1. For each genome, the optimal representative assembly was identified using <u>dRep</u> (13) v3.4.5. *A. johnsonii* E154408A strain was sequenced using second-generation DNA sequencing MiSeq platform (PE150; Illumina, Inc., San Diego, California, US). Raw reads were filtered using <u>fastp</u> (14) v0.23.4 using default settings. Filtered 1,322,156 reads were retained for genome assembly, conducted using <u>Shovill</u> v1.1.0 (**Table S1**).

### In vitro and in silico species typing

To provide a broader genomic perspective of the 22 PHSF-derived isolates and E154408A strain, an Average Nucleotide Identity (ANI) cluster map was generated using <u>ANIclustermap</u> including all NCBI RefSeq- and GenBank-available *A. johnsonii* genomes (n=89) excluding metagenome-assembled genomes (n=143) and anomalous assemblies (n=21) (**Table S2**). Among the PHSF-derived isolates, 3P2-tot-A was selected as a representative isolate based on the 99.9% ANI similarity observed across all PHSF-originating isolates and subjected to detailed genomic species typing. Confirmation of *A. johnsonii* 3P2-tot-A involved (i) ANI comparisons against *A. johnsonii* reference genome ANC 3681 (GenBank: GCA_000368805.1) and type strain CIP 64.6^T^ (GenBank: GCA_000368045.1); (ii) digital DNA–DNA hybridization (dDDH) using the <u>Genome-to-Genome Distance Calculator</u> (15) v3.0; and (iii) BLASTN analysis of *gyrB* and 16S rRNA gene against the *A. johnsonii* reference genome and type strain.

Due to the extended time gap between isolation (June 2007) and analysis (2024), only 14 of the original 22 PHSF-derived isolates could be successfully revived from the freezer stock for phenotypic species confirmation. In addition to the genome-based phylogeny, taxonomic placement of these recovered isolates, along with E154408A, underwent purity and species confirmatory testing using Matrix-Assisted Laser Desorption/Ionization Time-of-Flight (MALDI-ToF) mass spectrometry (Bruker microflex, Bruker Daltonics GmbH, Bremen, Germany) (16).

### Genome annotation and phylogenomic analysis

Assembled genomes were assessed for quality, completeness and contamination using <u>quast</u> v5.3.0 and <u>checkM</u> (17) v1.2.3 respectively. All assembled genomes were annotated using <u>Prokka</u> (18) v1.14.5. A total of 112 *A. johnsonii* genomes were included in all subsequent analyses, distributed as follows: (i) 89 genomes from NCBI RefSeq and GenBank (all available *A. johnsonii* genomes excluding metagenome-assembled and atypical genomes); (ii) 22 PHSF-derived isolates, and (iii) E154408A. Utilizing these genomes, <u>Roary</u> (19) v3.13.0 was used to generate *A. johnsonii* pangenome and define the species core genome.

A Maximum Likelihood phylogeny of the PHSF-derived isolates and E154408A strain was constructed based on a multi-sequence alignment of all core genes (n=976) using <u>RAxML</u> (20) v8.2.13 with GTRGAMMA model and 1,000 bootstraps. The reference genome of the taxonomically adjacent *Acinetobacter haemolyticus* HW-2A (GenBank: GCA_003323815.1) served as an outgroup in the phylogenomic analysis. The <u>ETE toolkit</u> (21) was used for tree manipulation, analysis, and visualization. A SNP analysis was conducted to assess clonality of PHSF-originating isolates using <u>snipit</u>, selecting *A. johnsonii* 2P07AA as the earliest reference genome.

### Functional annotation of nucleic acid sequences and metabolic mapping

Functional annotation of nucleic acid sequences from all genomes of *A. johnsonii* (n=112) according to KEGG Orthology, Enzyme Commission (EC), and COG category annotations was performed with <u>EggNOG-mapper</u> (22) v2.1.12 using MMseqs for the search step. Resulting annotated genes were subsequently represented in 232 metabolic maps using <u>KEGGCharter</u> (23) v1.1.2.

### Species-wide characterization of the resistome, putative virulence associated genes, plasmidome, and antibacterial biocide and metal resistance genes profile of *A. johnsonii*

All 112 available genomes of *A. johnsonii*, including PHSF-originating isolates and E154408A strain, were screened for the presence of i) ARGs; ii) virulence factors (VFs); and iii) antibacterial biocide- and metal-resistance genes using <u>ABRicate</u> v1.0.0 with the CARD 2023 (24), VFDB 2022 (25), and Bacmet (26) v2.0 databases, respectively. Only hits showing minimum 80% sequence identity and 60% coverage were retained. A custom database containing all *ade* homologues across *Acinetobacter* species was used to search for Ade efflux-pump-encoding genes. Species-wide plasmidome was characterized using <u>MOB-suite</u> (27) v3.1.9 with the sub-module MOB-recon v3.1.9. <u>ABRicate</u> v1.0.0 was ultimately used on the predicted plasmid sequences to identify plasmidic ARGs.

### Antimicrobial Susceptibility Testing (AST)

AST was performed on the available PHSF-originating isolates (n=14) and E154408A strain. 12 clinically relevant antimicrobials with interpretive criteria under EUCAST and CLSI guidelines were selected for testing: Piperacillin (100 μg), Ampicillin-sulbactam (10/10 μg), Cefepime (30 μg), Cefotaxime (30 μg), Imipenem (10 μg), Meropenem (10 μg), Gentamicin (10 μg), Tobramycin (10 μg), Amikacin (30 μg), Doxycycline (30 μg), Tetracycline (30 μg) and Ciprofloxacin (5 μg). Quality control strains *Escherichia coli* ATCC 25922 and *Pseudomonas aeruginosa* ATCC 27853 were included and maintained at standard conditions (35°C ± 2°C) according to the CLSI and EUCAST guidelines. However, as optimal growth for *A. johnsonii* occurs at temperatures ranging from 15 to 30°C (28), *A. johnsonii* isolates were incubated at 28-30°C. Clinical breakpoints for *Acinetobacter* species defined and established by the CLSI guidelines (CLSI M100-ED34:2024) were applied.

### Estimating the abundance of *A. johnsonii* in NASA cleanrooms

Raw shotgun metagenomic reads from two NASA cleanrooms, namely SAF at Jet Propulsion Laboratory (JPL, California, USA) and PHSF at KSC and two time points (i.e., 2016 and 2018) were acquired from the NCBI-SRA database (NCBI Project Accessions: PRJNA1150505 and PRJNA641079). Particularly, the metagenomic reads from 2016 were obtained from SAF at JPL (29), whereas the reads from 2018 were obtained from both cleanrooms (30). The metagenomic data from 2016 included 236 samples, of which 116 were treated with propidium monoazide (PMA) to selectively remove genomic DNA of dead bacterial targets (31). Data from 2018 included 94 samples, of which 24 PMA-treated and 23 PMA-untreated were from JPL-SAF and KSC-PHSF, respectively. Quality control and data filtering of raw metagenomic reads was performed using <u>fastp</u> (14) v0.22.0. <u>MetaCompass</u> v2.0 (32) was used to map filtered metagenomic reads from 2016 and 2018 to *A. johnsonii* 2P07AA, selected as the earliest reference genome, to gain insights into the abundance and prevalence of *A. johnsonii* in JPL and KSC controlled environments. The fraction of mapped reads to the reference genome was used to calculate the corresponding breadth of coverage for PMA-treated and untreated samples from both 2016 and 2018, setting a contig length cut-off score of 1000 bp.

## Results

### Species-wide ANI comparison

A species-wide all-vs-all ANI comparison of *A. johnsonii* revealed a distinct cluster of the 22 PHSF-derived isolates, each sharing 99.9% ANI (**Figure S1**). In contrast, PHSF-derived isolates exhibited ANI values ranging from 95.80% to 95.89% when compared to E154408A, confirming same-species relatedness but with notable genomic divergence. Based on the over 99.9% ANI based relatedness between PHSF-originating genomes, 3P2-tot-A isolate was selected as representative for in-depth genomic species typing of the PHSF-derived cluster, alongside E154408A.

### *In silico* species typing

Over 99% 16S rRNA and *gyr*B sequence identity of 3P2-tot-A isolate to both *A. johnsonii* reference genome ANC 3681 (95.90%) and type strain CIP 64.6^T^ supports species-level identity (**Table 1**). Observed over 95% ANI between *A. johnsonii* 3P2-tot-A isolate to both ANC 3681 (95.90%) and CIP 64.6^T^ (95.83%) strains confirmed same-species relatedness (**Table 1**). >65% dDDH values between 3P2-tot-A isolate and both ANC 3681 (65.60%) and CIP 64.6^T^ (66.20%) strains further suggested species-level identity (33). Analogous results were obtained for E154408A strain (**Table 1**), confirming that both the PHSF-derived isolates and E154408A strain taxonomically belong to *A. johnsonii*.

**Table 1:** species typing of strains 3P2-tot-A and E154408A based on ANI, dDDH, 16S rRNA gene and *gyrB* sequence similarity compared to both *A. johnsonii* reference genome ANC 3681 and *A. johnsonii* type strain CIP 64.4.

| Comparison: | ANI | dDDH | 16S rRNA sequence identity | *gyrB* sequence identity |
| --- | --- | --- | --- | --- |
| *A. johnsonii* 3P2-tot-A vs ref. genome ANC 3681 | 95.5% | 65.60% | <99.74% | % 98.51 |
| *A. johnsonii* 3P2-tot-A vs type strain CIP 64.4 | 95.83% | 66.20% | 99.67% | % 97.21 |
| *A. johnsonii* E154408A vs ref. genome ANC 3681 | 95.92% | 66.10% | 99.58% | % 97.09 |
| *A. johnsonii* E154408A vs type strain CIP 64.4 | 95.81% | 65.70% | 99.65% | % 97.13 |

### Phylogenomic analysis

The phylogeny of the 22 *A. johnsonii* PHSF-derived isolates and E154408A strain was inferred using a core-genome based Maximum-Likelihood approach based on all available *A. johnsonii* genomes (n=112) (**Figure 1)**. The PHSF-derived isolates formed a distinct monophyletic clade differing by 40-77 core SNPs, diverging from the main species lineage. *A. johnsonii* M19 strain (environmental source, Shandong, China) was identified as the closest phylogenetic relative to this cluster, followed by a monophyletic clade comprising *A. johnsonii* strains C4 (environmental source, Cahuita National Park, Costa Rica), JH7 (environmental source, Guangxi, China), GD03955, and GD03761 (both from environmental sources, Pakistan). Meanwhile, *A. johnsonii* E154408A grouped with XY27 strain (animal source, Shanghai, China), GD03727 (environmental source, Pakistan), and mNGS2101_37 (human source, Hangzhou, China).

**Figure 1:**
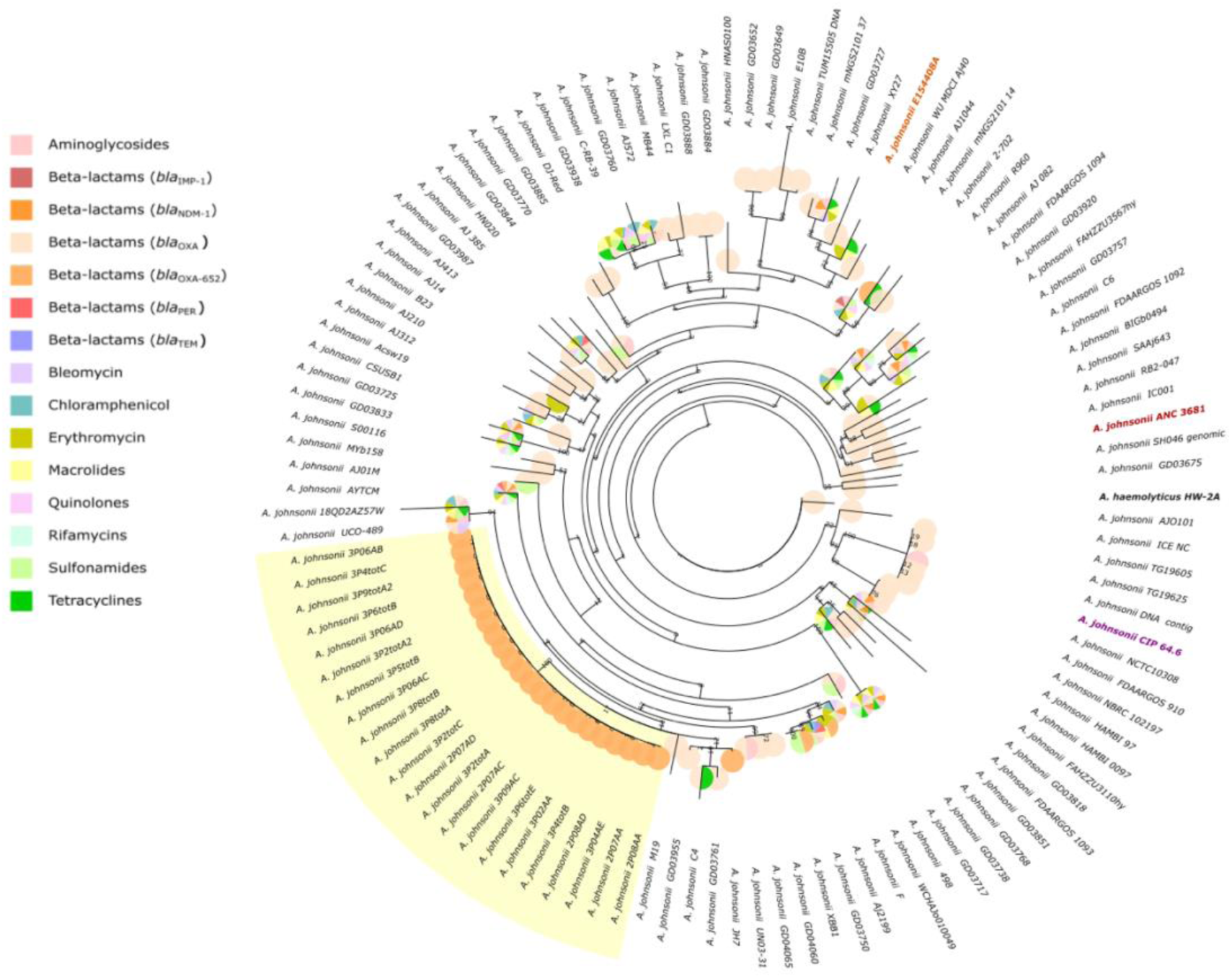
Phylogenomic analysis. Phylogeny of PHSF-originating isolates (highlighted in yellow) and E154408A strain (orange) inferred through Maximum Likelihood. Pie Charts associated with the terminal branches represent identified ARGs in correspondent genomes. *A. johnsonii* reference genome ANC 3681 and CIP 64.6^T^ type strain are highlighted in respectively red and purple. The reference genome of *A. haemolyticus* HW-2A served as an outgroup.

### Functional annotation of nucleic acid sequences and metabolic mapping

A total of 3,147 different genes across all *A. johnsonii* genomes (n=112) were functionally annotated using eggNOG-mapper. An average of 2,070 (range: 2,052-2,075) genes were annotated in PHSF-originating isolates, one of these coding for an allantoin racemase (KO: K16841; EC: 5.1.99.3; COG4126) involved in purine metabolism which was exclusively detected in PHSF-originating genomes except for 2P08AD strain (**Figure 2A**). 2,050 genes were annotated in the genome of *A. johnsonii* E154408A, one of these coding for an alkylglycerone-phosphate synthase (K00803; EC number 2.5.1.26; COG0277) which was not detected in any other *A. johnsonii* genome (**Figure 2B**).

**Figure 2:**
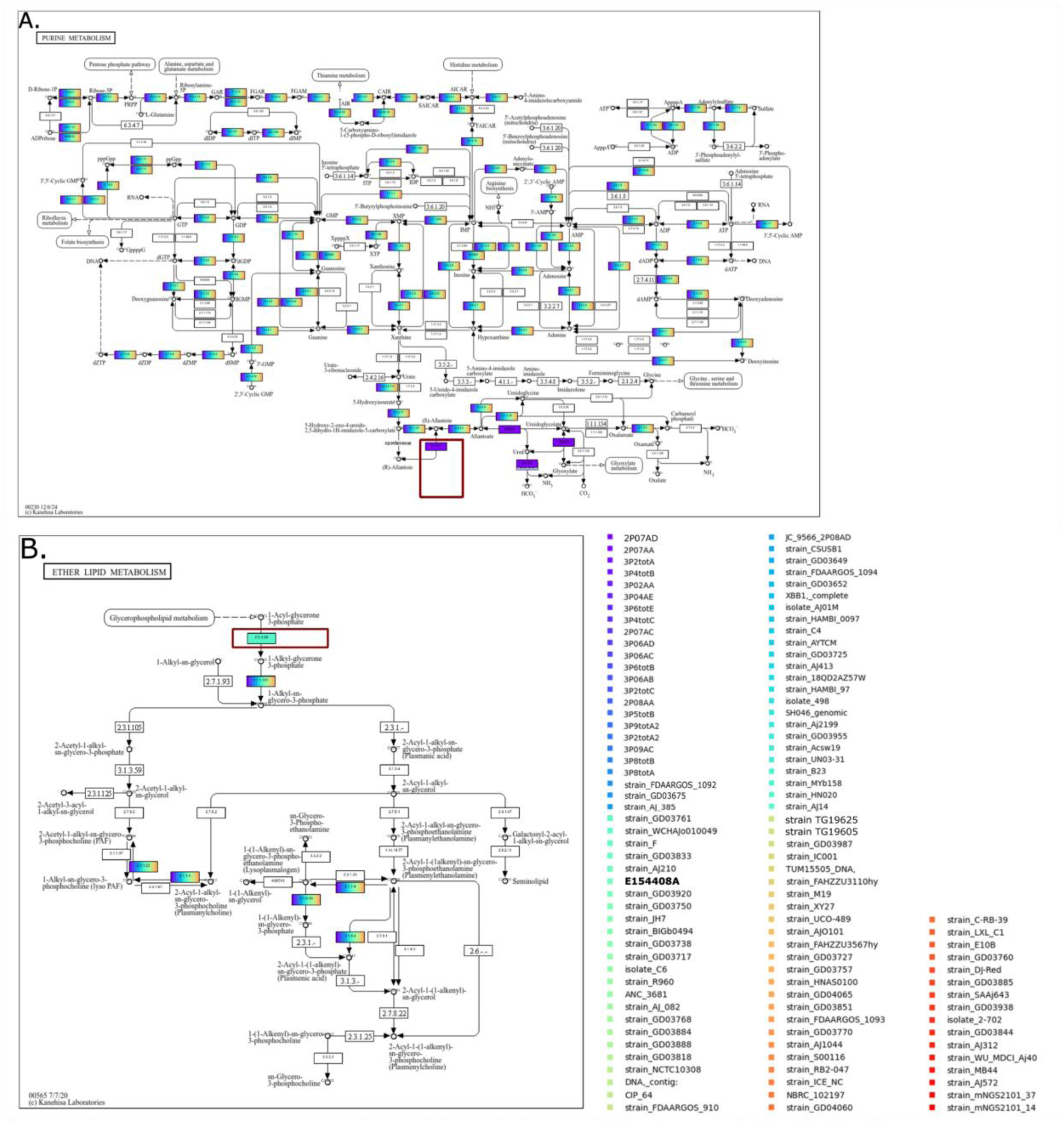
Functional annotation of nucleic acid sequences and metabolic mapping. Genome-based metabolic profiling and pathway predictions represented using KEGGCharter. **(A)** PHSF-derived isolates encode a unique allantoin racemase (red square) involved in purine metabolism; **(B)** E154408A strain encodes an alkylglycerone-phosphate synthase involved in ether lipid metabolism which was not detected in any other *A. johnsonii* genome.

### Species-wide characterization of the plasmidome of *A. johnsonii*

A total of 89 distinct plasmids were predicted across all *A. johnsonii* genomes, including 33 newly described types. Of these, 51 were assigned a relaxase (57.30%) (**Table S3**). Genomes were identified to carry between 0 and 8 putative plasmids (average per genome 1.83). AC600 was predicted with the highest frequency across the dataset, appearing in 19/112 (16.96%) of the genomes, followed by AC961 (14/112, 12.50%). While 5 PHSF isolates lacked detectable plasmids, the rest of the PHSF-derived isolates (n=17) consistently encoded a shared set of 2 putative plasmids (AD642 and AC600) predicted to be originating from *Acinetobacter wuhouensis* (34) and *A. johnsonii*, respectively. Notably, AC600 was found in 15/22 (68.19%) of the PHSF-derived *A. johnsonii* genomes and only four other *A. johnsonii* genomes from the broader dataset. Based on Mash distance neighbor analysis, AC600 is highly similar to a plasmid originating from isolate *A. johnsonii* XBB1 (mash distance 0.005). Two additional plasmids (AH350 and AD731), respectively predicted to be originating from *Moraxella osloensis* (35) and *A. baumannii* (36), were identified in respectively 2P07AA and 3P06AC strains. The clinical strain E154408A encoded six distinct putative plasmid types, including AH350 previously found in one PHSF-originating isolate.

### Antimicrobial resistance

A comprehensive search for ARGs in all available genomes of *A. johnsonii* led to the identification of a species-wide resistome comprising 62 ARGs predicted to confer resistance to aminoglycosides (*aac*(3), *aac*(6’), *ant*(2’’), *aph*(3’), *aph*(4), *aph*(6)-type genes), carbapenems (*bla*_IMP-1_, *bla*_OXA_, *bla*_NDM-1_, *bla*_PER-1_, *bla*_PER-2_, *bla*_TEM-1_, *BRP(MBL)*), macrolides (*ereA2*, *estT*, *mphE*, and *msrE*), phenicols (*catB3*, *cmlB1, floR*), quinolones (*qnrVC6*), sulfonamide (*sul1* and *sul2*), tetracycline (tet genes), and rifamycin *(arr-3)* (**Figure 3**). Among these, 37 ARGs were identified on predicted plasmid sequences in 35 genomes of *A. johnsonii* **(Table S4)**. *A. johnsonii* AYTCM strain (human source, Zhejiang, China) was observed to encode the highest number of ARGs (n=19), including multiple genes conferring resistance to carbapenems, namely *BRP(MBL)*, *bla*_IMP-1_, *bla*_NDM-1_, *bla*_OXA-58_, *bla*_OXA-652_, *bla*_PER-1_. Of note, 100% (112/112) of *A. johnsonii* genomes were observed to carry at least one *bla*_OXA_. One single ARG, *bla*_OXA-652_, potentially conferring broad-spectrum resistance to ß-lactams, was chromosomally detected in all PHSF-originating *A. johnsonii* genomes. *bla*_OXA-652_ was also detected in *A. johnsonii* strains mNGS2101_14 (human source, Hangzhou, China), JH7 (environmental source, Guangxi, China), GD03750 (environmental source, Pakistan), F (environmental source, Taizhou, China), AYTCM (human source, Zhejiang, China), Acsw19 (environmental source, Luzhou, China), and XBB1 (environmental source, Chengdu, China). A total of 5 ARGs were chromosomally detected in *A. johnsonii* E154408A strain, namely *bla*_OXA-212_, *mphE*, *msrE*, and *tet*(*39*), potentially conferring resistance to ß-lactams, macrolides, sulfonamide, and tetracycline antibiotics. However, these ARGs were found in a variable number (range: 8-26) of other *A. johnsonii* genomes. Of note, a second oxacillinase gene, *bla*_OXA-23_, was detected in E154408A and predicted to have plasmidic location (plasmid AD622). *bla*_OXA-23_ was also predicted on the same plasmid in *A. johnsonii* M19 strain.

**Figure 3:**
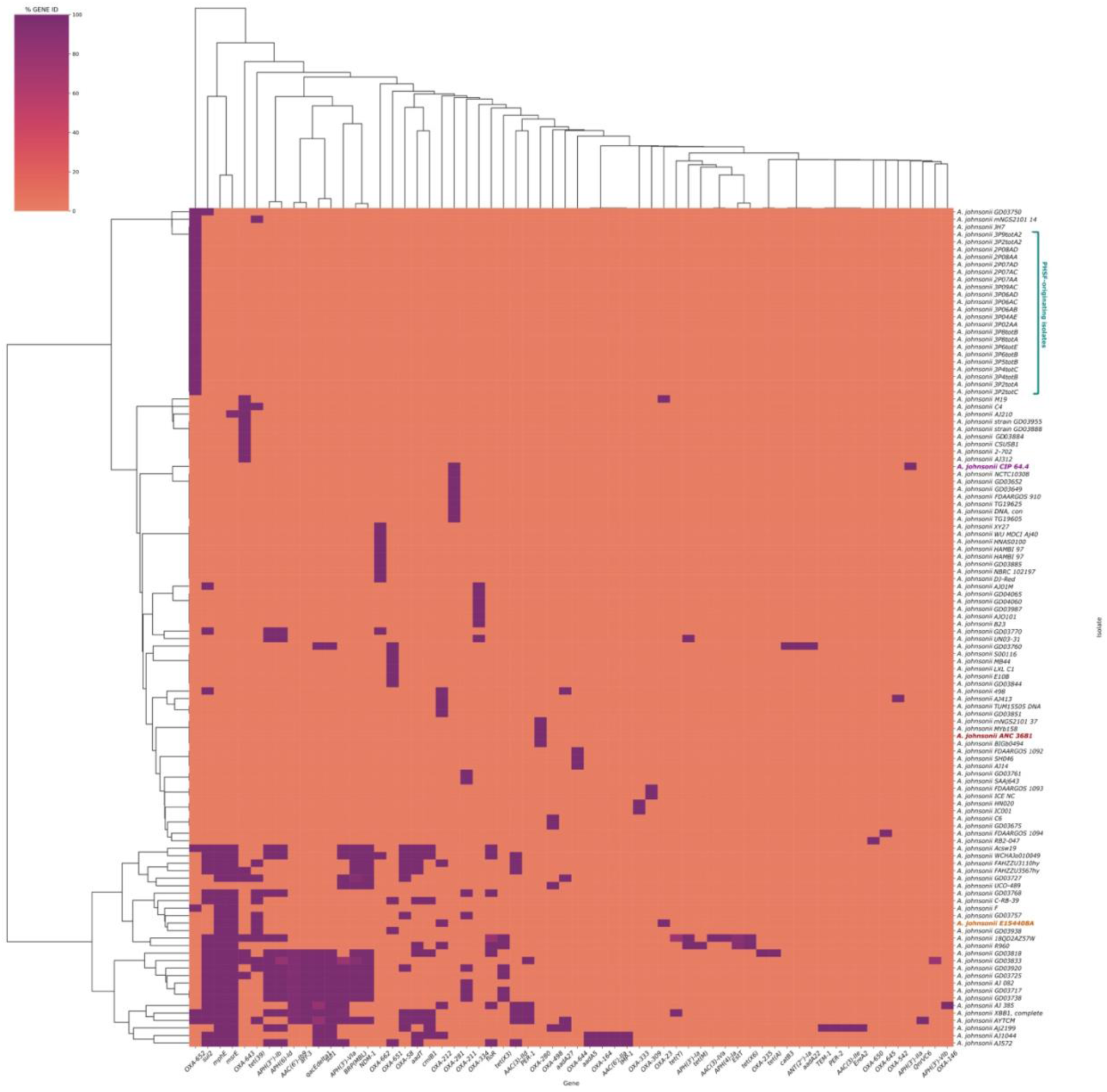
Species-wide characterization of the resistome of *A. johnsonii*. Species-wide characterization of the resistome of *A. johnsonii* (n= 112 strains), including PHSF-originating genomes (teal) and *A. johnsonii* E154408A (orange). *A. johnsonii* reference genome and type strain are highlighted in respectively red and purple.

Phenotypic AST revealed that nine out of the 14 tested PHSF-originating isolates (i.e., E154408A, 3P2-tot-C, 3P5-tot-B, 3P6-tot-E, 3P8-tot-B, 3P04AE, 3P06AC, 3P06AD, 3P09AC) showed intermediate susceptibility to cefotaxime, while the remaining isolates were sensitive (**Figure S4**). In addition, five isolates (i.e., 3P6-tot-E, 3P8-tot-B, 3P04AE, 3P06AD, 3P09AA) demonstrated intermediate susceptibility to piperacillin, with the rest being sensitive. Notably, *A. johnsonii* E154408A exhibited multi-drug phenotypic resistance to six out of the 12 tested antimicrobials, particularly to tetracycline, piperacillin, cefepime, and ciprofloxacin as well as the carbapenems imipenem and meropenem.

### Species-wide characterization of putative virulence associated genes of *A. johnsonii*

A total of eight known VFs were identified across all available *A. johnsonii* genomes (**Figure S3**). *pilT* and *pilG* were ubiquitously found in 100% (112/112) of the genomes and *ompA* in 98.2% (110/112) of the genomes. PHSF-originating *A. johnsonii* genomes were observed to carry the same subset of 6 VFs, *ompA*, *pilG*, *pilT*, *hcp/tssD*, *tssC*, and *tse4*. 3 VFs, *pilG*, *pilT*, and *ompA*, were identified in E154408A strain.

### Species-wide characterization of antibacterial biocide and metal resistance gene profiles in *A. johnsonii*

A total of 146 antibacterial biocide and metal resistance genes were identified across the dataset using the BacMet database (**Figure 4**). Highly prevalent genes, appearing in over 97% of the genomes used, included the AdeIJK efflux pump genes *adeI/J/K,* which mediate resistance to a broad range of antibiotics (including beta-lactams and fluoroquinolones) and anionic surfactant compounds; *oxyRkp* (resistance to hydrogen peroxide and detergents); *fabI* and *mexT* (triclosan); *sitB* and *sodB* (hydrogen peroxide); *rpoS* and *gadC/xasA* (hydrochloric acid); *evgA* (sodium deoxycholate); and multiple metal resistance genes.

**Figure 4:**
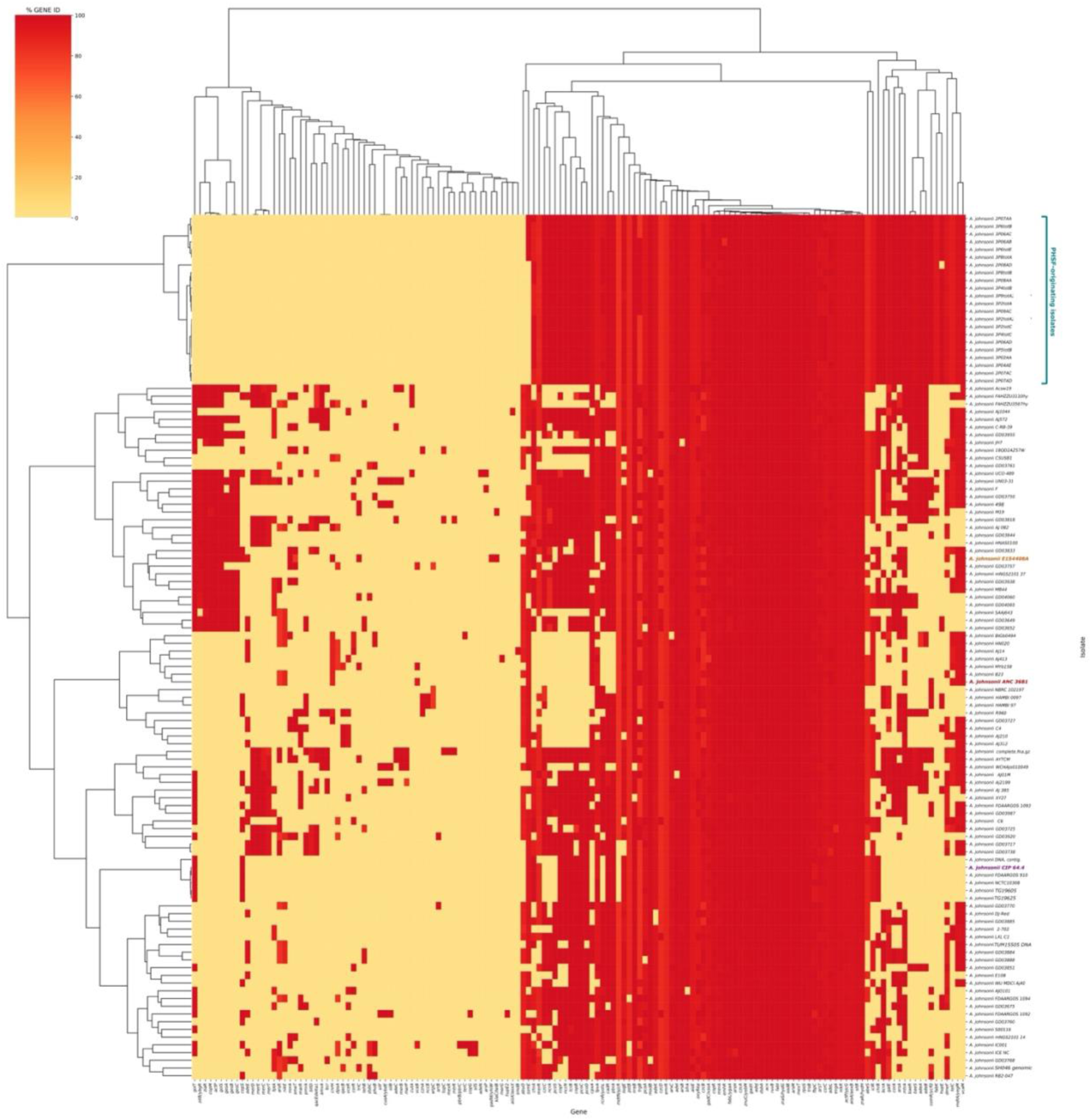
Species-wide characterization of antibacterial biocide and metal resistance gene profiles in *A. johnsonii*. Species-wide characterization of the profile of antibacterial biocide- and metal-resistance genes using BacMet in *A. johnsonii* (n= 112 strains), including PHSF-originating genomes (teal) and *A. johnsonii* E154408A (orange). *A. johnsonii* reference genome and type strain are highlighted in respectively red and purple. A total of 146 antibacterial biocide and metal resistance genes were identified across the dataset. Highly prevalent genes, appearing in over 97% of the genomes used, included *adeI/J/K,* which mediate resistance to a broad range of antibiotics (including beta-lactams and fluoroquinolones) and anionic surfactant compounds, *oxyRkp* (resistance to hydrogen peroxide and detergents); *fabI* and *mexT* (triclosan resistance); *sitB* and *sodB* (hydrogen peroxide); *rpoS* and *gadC/xasA* (hydrochloric acid); *evgA* (sodium deoxycholate); and multiple genes conferring resistance to arsenic, cadmium, cobalt, copper, iron, magnesium, mercury, and zinc. An average of 82 (range: 81–83) of the 146 identified genes were detected among the 22 PHSF-derived genomes while 87 were identified in the clinical E154408A strain. While PHSF-originating isolates and E154408A carried the complete AdeIJK efflux pump gene cassette, PHSF genome additionally possessed *adeA* and *adeB*, two components of the AdeABC multidrug resistance efflux pump also found in 43.75% of the rest of the dataset.

Among the 22 PHSF-derived genomes, an average of 82 (range: 81–83) of the 146 identified genes were detected. Notably, *ssmE*, encoding for the SsmE multidrug efflux pump, was present in only 5.36% (6/22) of the PHSF-derived genomes compared to 73.03% (65/112) of the remaining *A. johnsonii* isolates. 87 antibacterial biocide and metal resistance genes were identified in the clinical E154408A strain. While PHSF-originating isolates and E154408A carried the complete AdeIJK efflux pump gene cassette, PHSF genomes additionally possessed *adeA* and *adeB*, two components of the AdeABC multidrug resistance efflux pump also found in 43.75% of the rest of the dataset. Further screening with a custom database revealed the presence of *adeW* and *adeT* (homologues of *adeA*) and *adeR* and *adeS* (other components of AdeABC) in all PHSF isolates, though *adeS* showed reduced sequence identity relative to the reference sequence. The *adeN* repressor of AdeIJK was also detected in all PHSF-derived isolates, albeit with diminished identity to the reference.

### Estimating the abundance of *A. johnsonii* in NASA cleanrooms

Metagenome mapping with MetaCompass was conducted to estimate the prevalence of *A. johnsonii* in NASA cleanrooms (**Figure 5A-C**). In samples from JPL-SAF, 2016, the largest percentage of mapped reads obtained was 6.62% for PMA-untreated and 5.37% for PMA-treated samples, suggesting overall low yet persisting presence of either viable cells or residual genomic signatures of *A. johnsonii* in the cleanroom. While breadth of coverage of *A. johnsonii* genome in PMA-treated samples was generally low (average 12.51%) across all locations, higher values (average 27.88%) were measured in PMA-untreated samples with a peak of 76.68% at location 11 (**Figure 5D**). A maximum value of 1.92% mapped reads was obtained in PMA-treated samples from JPL-SAF, 2018, suggesting the persistence of a minimal population of viable cells of *A. johnsonii* in the cleanroom (**Figure 5E**). However, up to 9.05% reads were mapped in PMA-untreated samples, indicating a significant fraction of the signal coming from non-viable cells. Analogously, low breadth of coverage (average 4.15; maximum 5.99%) was measured in PMA-treated samples, whereas up to 71.68% coverage (average 25.87%) was obtained for PMA-untreated samples. Similarly, metagenome mapping with samples collected from KSC-PHSF in 2018 led to up to 1.87% mapped reads with low breadth of coverage (average 4.58%; maximum 11.03%) in PMA-treated samples, and up to 13.41% mapped reads with higher coverage (average 79.86%; maximum 89.37%) in PMA-untreated samples (**Figure 5F**). Collectively, these results suggest a higher *A. johnsonii* load in the samples collected from KSC-PHSF in 2018 compared to those collected from JPL-SAF in the same year.

**Figure 5:**
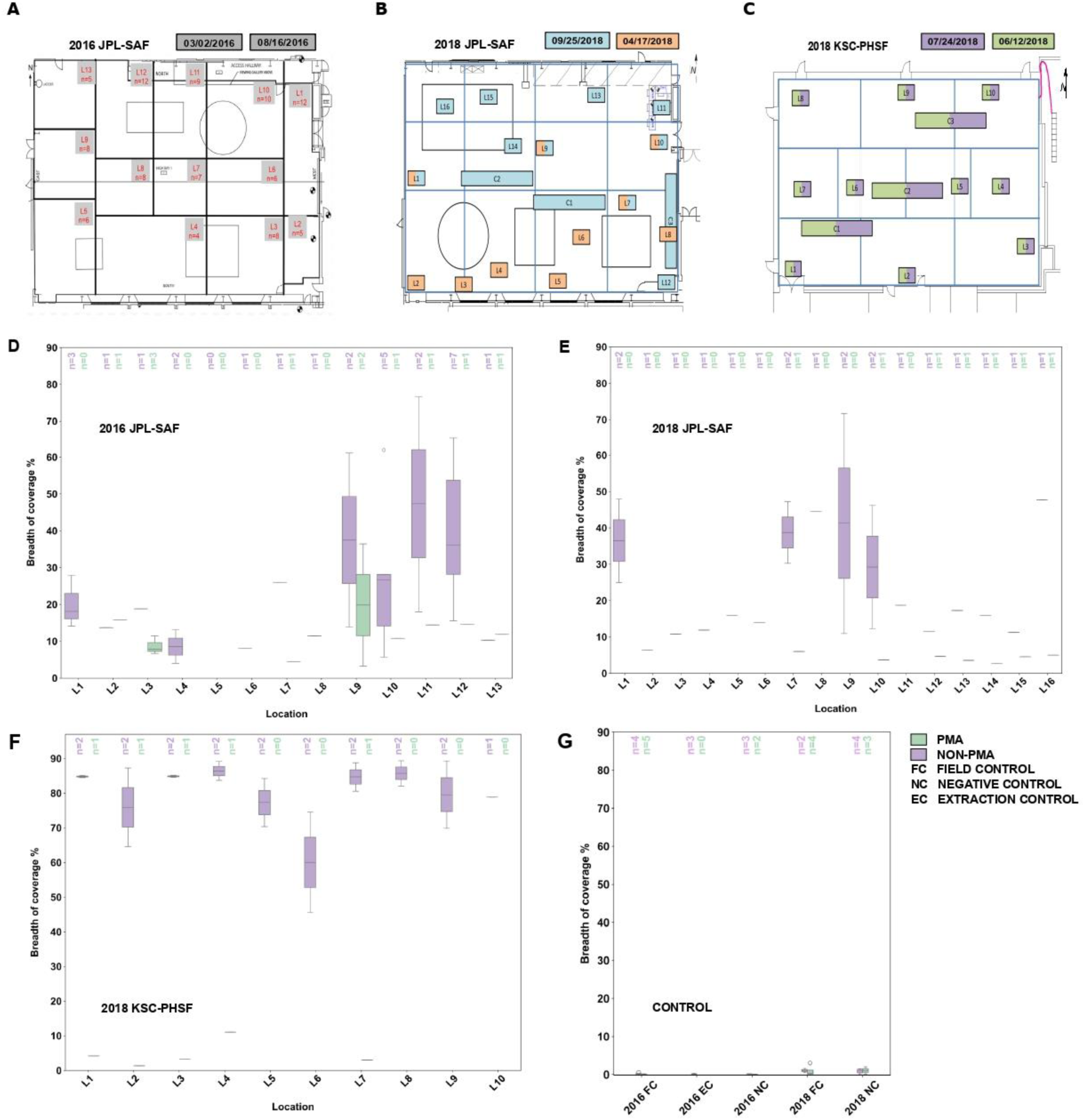
Estimating the abundance of *A. johnsonii* in NASA cleanrooms. Metagenome mapping results of *A. johnsonii* in cleanrooms. **(A-C)** Schematic representation of dates and specific locations for samples collected at JPL-SAF over a six-month period in 2016 **(A)**; at JPL-SAF during two separate events in 2018 **(B)**; and at KSC-PHSF during two separate events in 2018 **(C)**. The colors of the squares correspond to the sampling date. The graph is divided into artificial quadrants based on sample grouping and foot traffic. Dates are presented in the format: Month/Date/Year; **(D-F)** Box plots showing the breadth of coverage in percentile for *A. johnsonii* for PMA-treated (PMA) vs PMA-untreated (non-PMA) across different sampling locations in 2016 JPL-SAF **(D)**; 2018 JPL-SAF **(E)**; and 2018 KSC-PHSF **(F)**; **(G)** Box plot depicting the breadth of coverage in percentile for controls, comparing PMA vs. non-PMA for years 2016 and 2018.

## Discussion

Taxonomic classification of both PHSF-originating isolates (n=22) and the clinical strain E154408A as *A. johnsonii* was confirmed through a combination of *in vitro* and *in silico* approaches, using both *A. johnsonii* reference genome and type strain CIP 64.6^T^ for comparison. Bootstrap-supported core pangenome–based phylogeny identified that the PHSF-derived isolates form a monophyletic clade diverging from the main *A. johnsonii* lineage, suggesting the potential emergence of genomic signatures favorable for adaptation to extremely clean environments.

While observed 40-77 core SNPs variation between the 22 PHSF-derived isolates is compatible with a single clonal lineage, this exceeds the threshold typically used to define outbreak-related strains in *Acinetobacter baumannii* (≤10 SNPs) (37), which is consistent with a dataset of closely related yet genetically distinct isolates.

Genome-based metabolic profiling and pathway predictions revealed only subtle differences between PHSF-originating and the remaining *A. johnsonii* genomes, indicating the absence of broad specialized pathways conferring adaptation to cleanroom conditions. Nonetheless, the PHSF-derived isolates uniquely encode an allantoin racemase gene absent in all other *A. johnsonii* genomes examined. Allantoin racemase catalyzes the interconversion of R- and S-allantoin, potentially enhancing the breakdown of purines into metabolically useful intermediates (38). Enhanced purine utilization could offer a nutritional edge in nutrient-poor environments by providing an additional route for nitrogen and carbon acquisition (38). Further research is needed to determine whether this unique and highly conserved trait contributes to more efficient nutrient utilization and, ultimately, to the survival of *A. johnsonii* in the cleanroom environment.

The number of predicted plasmids per PHSF-derived genome varied from zero to three, suggesting dynamic events of plasmid gain and/or loss across the monophyletic population and further attesting to the genetic diversification of PHSF-originating isolates. Notably, the nearest neighbor of one of the two mobilizable plasmids identified in E154408A (AC935) is known to carry *bla*OXA genes (39), indicating a potential pathway for the acquisition of *bla*OXA-23.

Due to the contig-level assembly status of these genomes, it remains challenging to definitively ascertain whether resistance and putative virulence-associated genes identified in these isolates are plasmid-encoded or located on chromosomal regions. Further high-resolution sequencing and functional studies would be necessary to clarify their genomic origins.

Species-wide characterization of the *A. johnsonii* resistome revealed the intrinsic presence of *bla*OXA in all examined genomes, with no exceptions. This is consistent with previous reports that *A. johnsonii* can persist in polluted environments containing antibiotic residues^4^ and highlight the risk posed by *Acinetobacter* species in causing difficult-to-treat infections due to limited therapeutic options (40). While the species-wide presence of *bla*OXA in every *A. johnsonii* genome is not common knowledge, our findings provide a large-scale confirmation of *bla*OXA ubiquity in *A. johnsonii*.

All PHSF-originating isolates consistently carried a single chromosomal ARG, namely *bla*OXA-652 from the *bla*OXA-211 family, known for its narrow spectrum cephalosporin hydrolysis and lack of clinically significant carbapenem resistance (41). Indeed, none of the PHSF-originating isolates exhibited carbapenem resistance at the phenotypic level. In contrast, E154408A displayed phenotypic resistance to carbapenems (imipenem and meropenem), cefepime, ciprofloxacin, piperacillin, and tetracyclines, consistent with its genomic profile marked by *mphE*, *msrE*, *tet*(*39*), and by the co-occurrence of the chromosomal *bla*OXA-212 and plasmidic *bla*OXA-23.

Although biosynthesis of these oxacillinases was not directly verified, the observed carbapenem resistant phenotype can be explained by the presence of *bla*OXA-23 (40). However, resistance of E154408A strain to carbapenems can also be interpreted as an additive resistance phenotype due to the cumulative presence of intrinsic and acquired oxacillinases, which elevate antimicrobial resistance levels beyond what a single enzyme would achieve (42). Collectively, these findings highlight *A. johnsonii* as an understudied yet emerging antimicrobial resistance threat and suggest the potential ability of this species to develop enhanced antimicrobial resistant profiles due to the combined presence of chromosomal and acquired oxacillinases.

We highlight, however, the lack of clinically certified phenotypic AST assays tailored for *A. johnsonii*’s specific optimal growth conditions and of guidelines (e.g., CLSI and EUCAST) to interpret results accordingly. Addressing this gap is of importance to manage *A. johnsonii* with potential MDR phenotype in clinical setting decision-making.

The widespread presence of key components of the Type VI Secretion System (T6SS) (*tse4*, *tssC*, and *hcp/tssD*) among PHSF-deriving isolates, less commonly found in the broader dataset, aligns with evidence that a more complete T6SS pathway may assist their tolerance to nutrient-scarce environments. Additionally, the fact that PHSF-originating genomes belong to the subset of *A. johnsonii* strains encoding both AdeIJK and a nearly complete (missing only *adeC*) AdeABC efflux pumps may contribute to explain their resilience to repeated decontamination protocols, as supported by examples of increased tolerance of benzalkonium chloride, a sterilizing agent widely used in NASA cleanrooms, associated with the presence of AdeABC and AdeIJK efflux pumps in *A. johnsonii* (43). Notably, previous studies demonstrated the activity of AdeABC despite the absence of *adeC*, compensated by *adeK* (44).

Genome-to-metagenome mapping revealed the presence in PHSF of genomic signatures and a minimal population of viable cells of *A. johnsonii* over 10 years post initial isolation and overall greater bacterial load in KSC-PHSF compared to JPL-SAF, possibly due to the stricter disinfection regime applied in JPL-SAF following spacecraft assembly activities in 2018 (30).

In conclusion, our results suggest deviation of PHSF-derived isolates from the main *A. johnsonii* species lineage and the emergence in their genomes of unique and conserved traits that may explain adaptation of this non-spore-forming microorganism to extremely clean and nutrient-scarce environments. Further research, however, is required to demonstrate whether these genomic traits contributed to the survival of *A. johnsonii* in NASA cleanrooms. Through the comprehensive genomic characterization of *A. johnsonii*’s resistome, our results provide a valuable, large-scale confirmation of intrinsic *bla*OXA ubiquity in this species. With the documentation of the first reported carbapenem-resistant patient colonization case in Ireland and Europe, our findings also shed light on the potential of *A. johnsonii* to develop enhanced antimicrobial resistant phenotypes and the risks related to the persistence of this microbial contaminant as reservoir and vector of AMR in controlled environments. Collectively, our findings will help evaluate and manage the contamination risk of *A. johnsonii* across clinical, terrestrial and extraterrestrial settings.

## supplemental figures titles and legends

**Figure S1**: **Species-wide ANI comparison.** Species-wide all-vs-all ANI comparison of all available genomes of *A. johnsonii*, PHSF-originating isolates (teal), and E154408A strain (orange). *A. johnsonii* reference genome and type strain are highlighted in respectively orange and purple.

**Figure S2: SNP analysis.** SNP analysis conducted on the 22 PHSF-derived isolates of *A. johnsonii* using 2P07AA strain as the earliest reference genome. PHSF-originating isolates form a tight cluster differing by 40–77 core SNPs, consistent with a dataset of closely related yet genetically distinct isolates.

**Figure S3**: **Species-wide characterization of putative virulence-associated genes of *A. johnsonii*.** Species-wide characterization of putative virulence-associated genes using VFDB of *A. johnsonii* (n= 112 strains), including PHSF-originating genomes (teal) and *A. johnsonii* E154408A (orange). *A. johnsonii* reference genome and type strain are highlighted in respectively red and purple. A total of eight known VFs were identified across all genomes. *pilT* and *pilG* were ubiquitously found in 100% (112/112) of the genomes and *ompA* was detected in 98.2% (110/112) of the genomes. *hcp/tssD* and *tssC* were detected in the same subset of 86.6% (97/112) of the genomes. *tssC* was individually detected in strains HNAS0100 (clinical source, Hunan, China) and GD03725 (environmental source, Pakistan). GD04060, GD04065 (both environmental source, US), and UN03-31 (human source, Shenzhen, China) strains were identified to encode the highest number of VFs (n=7). PHSF-originating *A. johnsonii* genomes were observed to carry the same subset of 6 VFs, namely *ompA*, *pilG*, *pilT*, *hcp/tssD*, *tssC*, and *tse4*. 3 VFs were identified in E154408A strain, namely *pilG*, *pilT*, and *ompA*.

**Figure S4: AST.** AST interpreted under EUCAST and CLSI guidelines to 12 clinically relevant antimicrobials including Piperacillin (PRL), Ampicillin-sulbactam (SAM), Cefepime (FEP), Cefotaxime (CTX), Imipenem (IPM), Meropenem (MEM), Gentamicin (GEN), Tobramycin (TOB), Amikacin (AK), Doxycycline (DO), Tetracycline (TET) and Ciprofloxacin (CIP). Strains E154408A, 3P2-tot-C, 3P5-tot-B, 3P6-tot-E, 3P8-tot-B, 3P04AE, 3P06AC, 3P06AD, 3P09AC showed intermediate susceptibility (highlighted yellow) to CTX. Five strains (i.e., 3P6-tot-E, 3P8-tot-B, 3P04AE, 3P06AD, 3P09AA) demonstrated intermediate susceptibility to piperacillin. *A. johnsonii* E154408A exhibited phenotypic resistance (highlighted in red) to tetracycline, piperacillin, cefepime, and ciprofloxacin as well as carbapenems imipenem and meropenem.

## Resource availability

· The genomes described in this project have been deposited at DDBJ/ENA/GenBank under the BioProject numbers PRJNA1128436 and PRJNA1209713.

## Acknowledgments

This work was supported by Prof. Alexandre Soares Rosado’s KAUST Baseline Grant (BAS/1/1096-01-01). Part of the research described in this publication was carried out at the Jet Propulsion Laboratory, California Institute of Technology, under a contract with National Aeronautics and Space Administration. This research was funded by a 2012 Space Biology NNH12ZTT001N grant no. 19-12829-26 under Task Order NNN13D111T award to K.V. A.T. is supported by the Environmental Protection Agency Research Programme 2021-2030 as Government of Ireland initiative funded by the Department of the Environment, Climate and Communication. P.S. is supported through the Prime Minister’s Research Fellowship from the Ministry of Education, Government of India. F.M. is supported by Taighde Éireann – Research Ireland under Grant number GOIPG/2023/4515. The funders had no role in study design, data collection and interpretation, the writing of the manuscript, or the decision to submit the work for publication.

We thank Patrick Driguez, Angel Angelov and Alexander Putra at KAUST Core Laboratories, King Abdullah University of Science and Technology (KAUST), Thuwal, Kingdom of Saudi Arabia for sequencing the PHSF originating isolates. We thank Tahira Jamil at Biological and Environmental Sciences and Engineering Division (BESE), King Abdullah University of Science and Technology (KAUST), Thuwal, Kingdom of Saudi Arabia for assisting with genome assembly. We thank staff of Galway Reference Laboratory Services, Galway University Hospital, Galway, Ireland for reporting and sequencing the clinical originating isolate.

## Author contributions

Conceptualization, G.M. and K.V.; Methodology, G.M. and K.V.; Software, A.T., G.M., V.V. and P.S.; Formal Analysis, A.T., G.M., A.O’C., V.V. and P.S.; Investigation, A.T., G.M., A.O’C., F.M., A.K., B.H.; Resources, G.M., C.C., A.S.R., K.R. and K.V.; Data Curation, A.T., G.M.,; Writing - Original Draft, A.T., A.O’C., V.V., P.S., A.K.,; Writing - Review & Editing, A.T., G.M., N.K.S. and K.V.; Visualization, A.T. and V.V.; Supervision, G.M., K.R. and K.V.; Project Administration, A.T., G.M. and K.V.; Funding Acquisition, G.M.

## Supplemental information index

**Document S1:** Figures S1-S4.

**Table S1: Genome assembly statistics.**

**Table S2: List of publicly available genomes used.** Excel file containing a list of publicly available genomes used (too large to fit in a PDF).

**Table S3: Predicted plasmid information.** Excel file containing predicted plasmid information including plasmid id, size (bp), replicon and relaxase types (too large to fit in a PDF).

**Table S4: Plasmidic ARGs.** Excel file containing the results of ARGs screening on predicted plasmid sequences (too large to fit in a PDF).

