## Supplementary material for "Plasmidome, resistome, and virulence-associated genes characterization of *Acinetobacter johnsonii* in NASA cleanrooms and a clinical setting": Figures S1-S4

**Figure S1:**

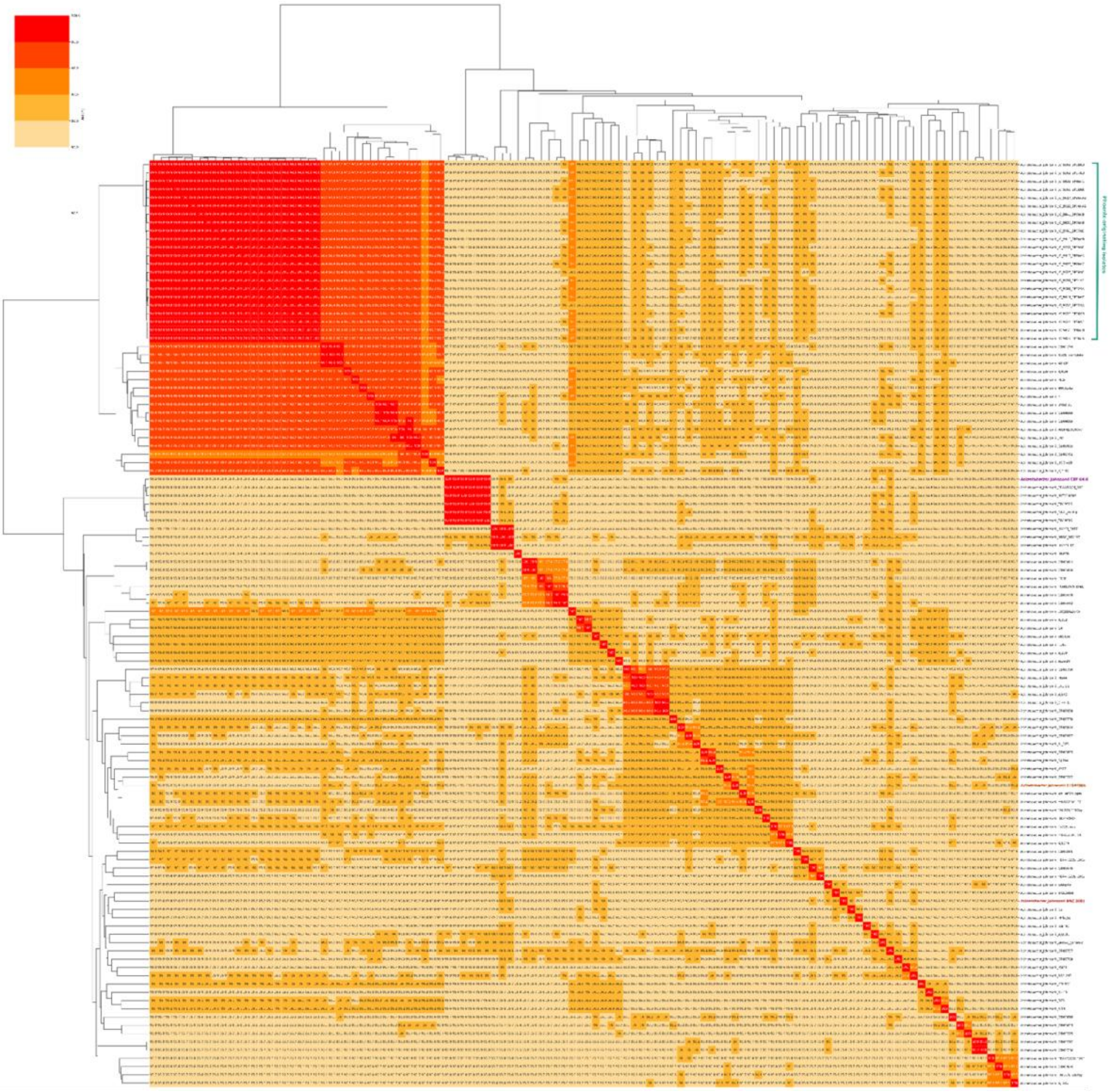

### Figure S2:

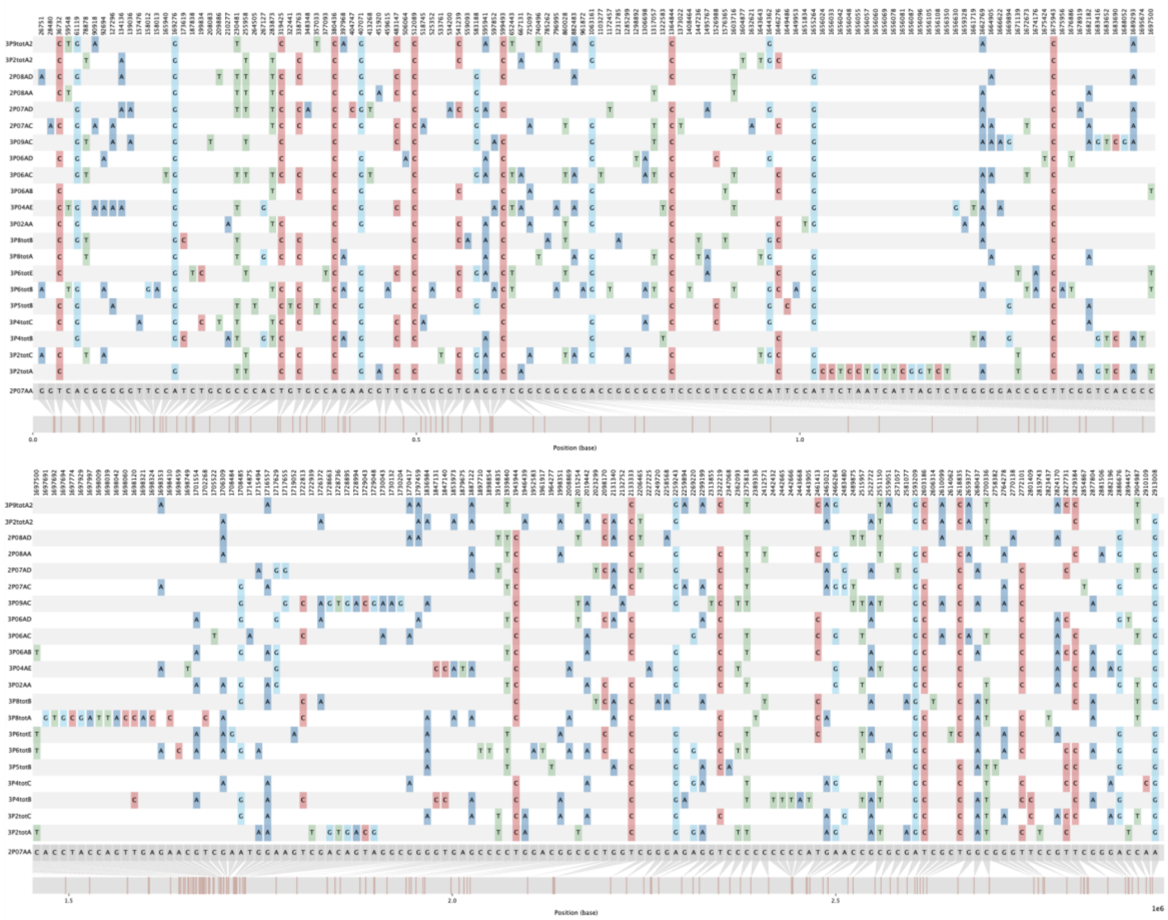

**Figure S3:**

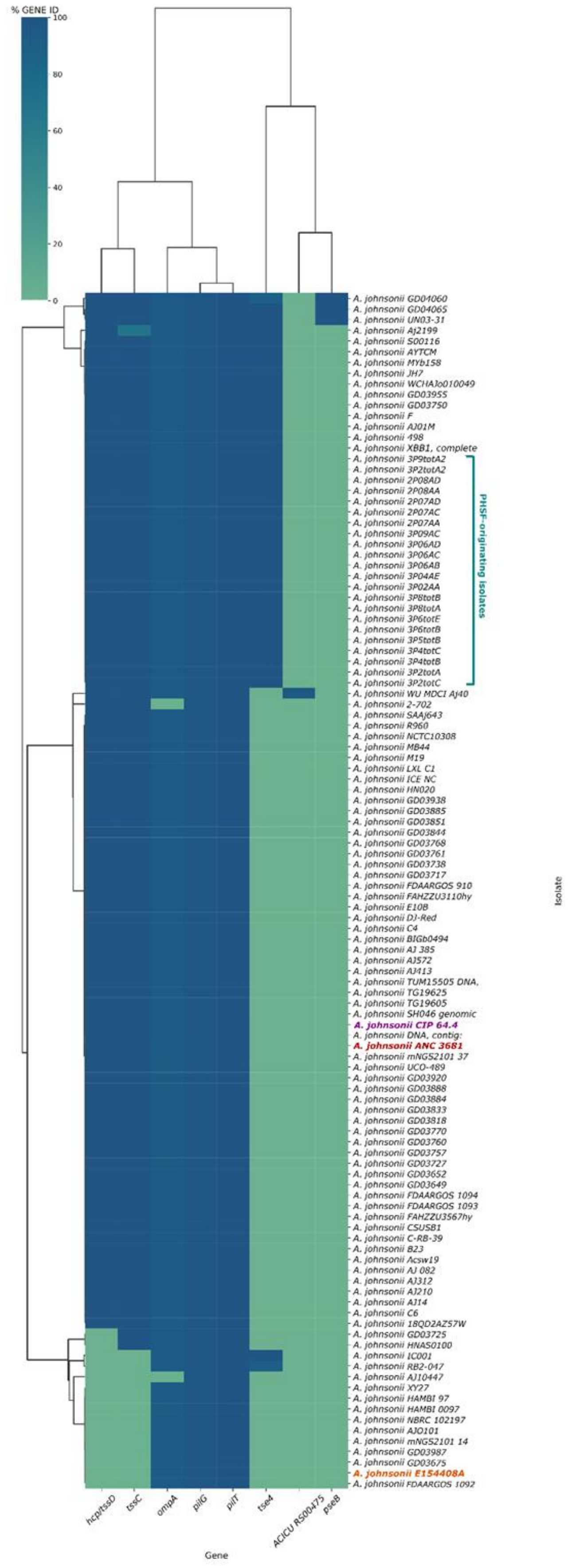
% GENE ID
100
80
60
40
20
0
A. johnsonii GD04060
A. johnsonii GD04065
A. johnsonii UN03-31
A. johnsonii Aj2199
A. johnsonii S00116
A. johnsonii AYTCM
A. johnsonii MYb158
A. johnsonii JH7
A. johnsonii WCHAJo010049
A. johnsonii GD03955
A. johnsonii GD03750
A. johnsonii F
A. johnsonii AJ01M
A. johnsonii 498
A. johnsonii XBB1, complete
A. johnsonii 3P9totA2
A. johnsonii 3P2totA2
A. johnsonii 2P08AD
A. johnsonii 2P08AA
A. johnsonii 2P07AD
A. johnsonii 2P07AC
A. johnsonii 2P07AA
A. johnsonii 3P09AC
A. johnsonii 3P06AD
A. johnsonii 3P06AC
A. johnsonii 3P06AB
A. johnsonii 3P04AE
A. johnsonii 3P02AA
A. johnsonii 3P8totB
A. johnsonii 3P8totA
A. johnsonii 3P6totE
A. johnsonii 3P6totB
A. johnsonii 3P5totB
A. johnsonii 3P4totC
A. johnsonii 3P4totB
A. johnsonii 3P2totA
A. johnsonii 3P2totC
PHSF-originating isolates
A. johnsonii WU MDCI Aj40
A. johnsonii 2-702
A. johnsonii SAAj643
A. johnsonii R960
A. johnsonii NCTC10308
A. johnsonii MB44
A. johnsonii M19
A. johnsonii LXL C1
A. johnsonii ICE NC
A. johnsonii HN020
A. johnsonii GD03938
A. johnsonii GD03885
A. johnsonii GD03851
A. johnsonii GD03844
A. johnsonii GD03768
A. johnsonii GD03761
A. johnsonii GD03738
A. johnsonii GD03717
A. johnsonii FDAARGOS 910
A. johnsonii FAHZZU3110hy
A. johnsonii E10B
A. johnsonii DJ-Red
A. johnsonii C4
A. johnsonii BIGb0494
A. johnsonii AJ 385
A. johnsonii AJ572
A. johnsonii AJ413
A. johnsonii TUM15505 DNA,
A. johnsonii TG19625
A. johnsonii TG19605
A. johnsonii SH046 genomic
A. johnsonii CIP 64.4
A. johnsonii DNA, contig:
A. johnsonii ANC 3681
A. johnsonii mNGS2101 37
A. johnsonii UCO-489
A. johnsonii GD03920
A. johnsonii GD03888
A. johnsonii GD03884
A. johnsonii GD03833
A. johnsonii GD03818
A. johnsonii GD03770
A. johnsonii GD03760
A. johnsonii GD03757
A. johnsonii GD03727
A. johnsonii GD03652
A. johnsonii GD03649
A. johnsonii FDAARGOS 1094
A. johnsonii FDAARGOS 1093
A. johnsonii FAHZZU3567hy
A. johnsonii CSUSB1
A. johnsonii C-RB-39
A. johnsonii B23
A. johnsonii Acsw19
A. johnsonii AJ 082
A. johnsonii AJ312
A. johnsonii AJ210
A. johnsonii AJ14
A. johnsonii C6
A. johnsonii 18QD2AZ57W
A. johnsonii GD03725
A. johnsonii HNAS0100
A. johnsonii IC001
A. johnsonii RB2-047
A. johnsonii AJ10447
A. johnsonii XY27
A. johnsonii HAMBI 97
A. johnsonii HAMBI 0097
A. johnsonii NBRC 102197
A. johnsonii AJO101
A. johnsonii mNGS2101 14
A. johnsonii GD03987
A. johnsonii GD03675
A. johnsonii E154408A
A. johnsonii FDAARGOS 1092
Isolate
hcp/tssD
tssC
ompA
pilG
pilT
tse4
ACICU RS00475
pseB
Gene

**Figure S4:**

| | TET | DO | AK | GEN | TOB | SAM | PRL | FEP | CTX | IPM | MEM | CIP |
|---|---|---|---|---|---|---|---|---|---|---|---|---|
| 3P2totA | 25(S) | 30(S) | 26(S) | 24(S) | 23(S) | 30(S) | 22(S) | 22(S) | 23(S) | 31(S) | 29(S) | 30(S) |
| 3P2totC | 26(S) | 29(S) | 26(S) | 23(S) | 23(S) | 33(S) | 22(S) | 22(S) | 22(I) | 32(S) | 30(S) | 30(S) |
| 3P4totC | 28(S) | 32(S) | 25(S) | 24(S) | 24(S) | 33(S) | 21(S) | 22(S) | 23(S) | 34(S) | 31(S) | 31(S) |
| 3P5totB | 27(S) | 29(S) | 26(S) | 23(S) | 23(S) | 35(S) | 21(S) | 22(S) | 22(I) | 31(S) | 29(S) | 30(S) |
| 3P6totB | 25(S) | 29(S) | 26(S) | 22(S) | 23(S) | 33(S) | 21(S) | 22(S) | 23(S) | 34(S) | 32(S) | 28(S) |
| 3P6totE | 27(S) | 31(S) | 25(S) | 23(S) | 23(S) | 31(S) | 20(I) | 21(S) | 22(I) | 33(S) | 30(S) | 29(S) |
| 3P8totA | 28(S) | 32(S) | 26(S) | 22(S) | 23(S) | 33(S) | 21(S) | 23(S) | 23(S) | 33(S) | 31(S) | 29(S) |
| 3P8totB | 28(S) | 32(S) | 26(S) | 24(S) | 25(S) | 31(S) | 20(I) | 21(S) | 21(I) | 34(S) | 30(S) | 29(S) |
| 3P02AA | 28(S) | 30(S) | 24(S) | 25(S) | 22(S) | 31(S) | 23(S) | 24(S) | 24(S) | 31(S) | 30(S) | 31(S) |
| 3P04AE | 27(S) | 31(S) | 26(S) | 22(S) | 23(S) | 31(S) | 20(I) | 20(S) | 21(I) | 32(S) | 30(S) | 28(S) |
| 3P06AB | 26(S) | 30(S) | 26(S) | 22(S) | 23(S) | 36(S) | 22(S) | 23(S) | 23(S) | 31(S) | 30(S) | 31(S) |
| 3P06AC | 28(S) | 32(S) | 26(S) | 24(S) | 23(S) | 32(S) | 21(S) | 21(S) | 22(I) | 33(S) | 29(S) | 28(S) |
| 3P06AD | 26(S) | 32(S) | 25(S) | 22(S) | 23(S) | 30(S) | 20(I) | 21(S) | 21(I) | 29(S) | 28(S) | 29(S) |
| 3P09AC | 25(S) | 29(S) | 26(S) | 23(S) | 23(S) | 31(S) | 22(S) | 21(S) | 22(I) | 32(S) | 28(S) | 28(S) |
| E154408A | 12(R) | 20(S) | 28(S) | 25(S) | 24(S) | 18(S) | 6(R) | 12(R) | 15(I) | 8(R) | 9(R) | 6(R) |
